# Ca_V_1.2 channelopathic mutations evoke diverse pathophysiological mechanisms

**DOI:** 10.1101/2022.06.13.495975

**Authors:** Moradeke A. Bamgboye, Kevin G. Herold, Daiana C.O. Vieira, Maria K. Traficante, Philippa J. Rogers, Manu Ben-Johny, Ivy E. Dick

## Abstract

The first pathogenic mutation in Ca_V_1.2 was identified in 2004 and was shown to cause a severe multisystem disorder known as Timothy syndrome (TS). The mutation was localized to the distal S6 region of the channel, a region known to play a major role in channel activation. TS patients suffer from life-threatening cardiac symptoms as well as significant neurodevelopmental deficits including autism spectrum disorder (ASD). Since this discovery, the number and variety of mutations identified in Ca_V_1.2 has grown tremendously, and the distal S6 regions remains a frequent locus for many of these mutations. While the majority of patients harboring these mutations exhibit cardiac symptoms which can be well explained by known pathogenic mechanisms, the same cannot be said for the ASD or neurodevelopmental phenotypes seen in some patients, indicating a gap in our understanding of the pathogenesis of Ca_V_1.2 channelopathies. Here, we use of whole cell patch clamp, quantitative Ca^2+^ imaging, and single channel recordings to expand the known mechanisms underlying the pathogenesis of Ca_V_1.2 channelopathies. Specifically, we find that mutations within the S6 region can exert independent and separable effects on activation, voltage-dependent inactivation (VDI) and Ca^2+^-dependent inactivation (CDI). Moreover, the mechanisms underlying the CDI effects of these mutations are varied and include altered channel opening and possible disruption of CDI transduction. Overall, these results provide a structure-function framework to conceptualize the role of S6 mutations in pathophysiology and offer insight into the biophysical defects associated with distinct clinical manifestations.

## Introduction

Ca_V_1.2 L-type Ca^2+^ channels are perhaps the most prevalent of the voltage-gated Ca^2+^ channels, existing in cardiac, neuronal and smooth muscle cells (1). Ca^2+^ entry through these channels is precisely controlled through multiple forms of regulation including voltage-dependent activation, voltage-dependent inactivation (VDI) and Ca^2+^-dependent inactivation (CDI) (2-6). Each of these regulatory processes are vital, and disruption of them is expected to produce adverse physiological consequences. Timothy syndrome (TS) represents one such class of mutations, in which a single point mutation within Ca_V_1.2 leads to a severe multisystem disorder characterized by developmental delays, autism spectrum disorder (ASD) and profound long-QT syndrome (LQTS) (7-9). The underlying cause of TS was first described in 2004 as a single point mutation (G406R) within the distal IS6 region of Ca_V_1.2 (8). This form of LQTS is among the most severe presentation of this syndrome, with patients exhibiting QT intervals between 480-730 ms (7, 8, 10, 11). As a result, patients suffer from life threatening arrhythmias, which are often fatal in early childhood. Furthermore, TS represents one of the most penetrant monogenic forms of ASD (7, 8, 12). Since the first discovery of the G406R mutation, a growing number of mutations have been identified in Ca_V_1.2, most of which result in significant cardiac phenotypes (7, 8, 13-15). However, only select mutations also cause neurological phenotypes including ASD or developmental delay (7, 8, 13, 15, 16). As the number and variety of identified Ca_V_1.2 mutations continues to expand, so to does the phenotypic variation observed in patients. In the heart, the importance of Ca_V_1.2 inactivation has long been recognized and there is a clear mechanistic link between excess Ca^2+^ entry through Ca_V_1.2 and prolongation of the cardiac action potential (AP) (17-19), the cellular hallmark for LQTS. However, the mechanistic link between Ca_V_1.2 mutations and neurological effects remains much less clear. Mutations within both Ca_V_1.2 and Ca_V_1.3 channels which cause large effects on channel gating have been linked to ASD, however some Ca_V_1.2 mutations with significant effects on channel gating produce no neurological phenotype, and simple gain-of-function/loss-of-function descriptions of Ca_V_1.2 mutations result in little correlation with patient phenotypes (13, 20, 21).

The original G406R TS mutation is located within the distal IS6 region of the channel (8), as is the G402S mutation which was identified soon after (7). Both mutations produce a marked deficit in VDI (7, 22), fitting with prior work in the Ca_V_1.3 channel which demonstrated that residues within the distal S6 region may be critical to the function of a ‘hinge-lid’ inactivation scheme (23). Moreover, the S6 is known to play a critical role in channel activation (23-26). It is therefore not surprising that each of these mutations alters the voltage dependence of channel activation, although in opposing directions (G406R, hyperpolarizing shift; G402S, depolarizing shift) (22). Finally, both mutations have been shown to cause a decrease in CDI, demonstrating multiple effects on channel gating. Interestingly, the effect on CDI for many mutations within this region of both Ca_V_1.2 (including G402S and G406R) and Ca_V_1.3 has been demonstrated to be a direct result of the change in channel activation (22, 23), implying that the S6 region of the channel may not play a direct role in CDI. While a growing number of Ca_V_1.2 mutations have been identified across multiple channel regions, a large subset of these pathogenic mutations appear within distal S6 regions of the channel. Thus, there may be common mechanistic elements which can be elucidated by evaluating the effects of these S6 mutations.

Here, we evaluate the biophysical impact of select Ca_V_1.2 mutations on CDI, VDI and channel activation, focusing primarily on mutations near the distal S6 region of the channel. Despite the locus of these mutations near the channel activation gate, we find that not all S6 mutations exert an effect on channel activation. In contrast to the canonical TS mutations, we find that the S6 mutations within this study have more selective effects on channel properties, sometimes altering CDI or VDI independently. Further investigation into the mechanisms underlying the gating changes in CDI-altering mutations reveals a remarkably diverse impact of S6 mutations. In particular, two mutations (I1166V and I1166T) caused a marked change in single channel properties indicative of enhanced entry into mode 2 gating, while another distal S6 mutation (I1475M) resulted in a selective CDI deficit, implicating a greater role for the IVS6 region in CDI. Finally, we show that these biophysical changes produce a difference in the response of the channels to a neurological stimulus, demonstrating that the neurological phenotype may be most strongly associated with enhanced channel activation.

## Results

### Pathogenic Ca_V_1.2 mutations often appear near the distal S6

The first TS mutations described (G406R and G402S) were both located within the IS6 region and caused changes to channel activation (7, 8, 22). Since this first description, numerous additional mutations in proximity of the distal S6 of Ca_v_1.2 have been reported (13, 14, 16, 27). Given the importance of this region in channel regulation, we chose several pathogenic mutations near a distal S6 region of the channel to focus on. **Fig. 1A** displays the locus of each mutation on a cartoon of the Ca_V_1.2 channel, with the S6 regions highlighted in blue for easy reference. Mutation L762F (14) resides just past the IIS6 region and has been shown to cause a cardiac selective phenotype. We next considered two mutations within the IIIS6 region. Interestingly I1166T and I1166V (13) occur at the identical residue, yet produce distinct phenotypes, with I1166T producing a multisystem disorder including neurological effects, while I1166V causes a cardiac selective effect. From the IVS6 domain we focused on the mutation I1475M, which has been described as cardiac selective in patients (13). Finally, we considered the effect of one mutation which resides outside an S6 region. E1496K resides within the EF-hand located on the C-tail of the channel and is associated with a cardiac-selective phenotype (13). This mutation is included in this analysis as this region has previously been identified as critical for VDI (26, 28), and has been suggested to cause a shift in channel activation (13) similar to the effects seen with S6 mutations.

**Figure 1.**
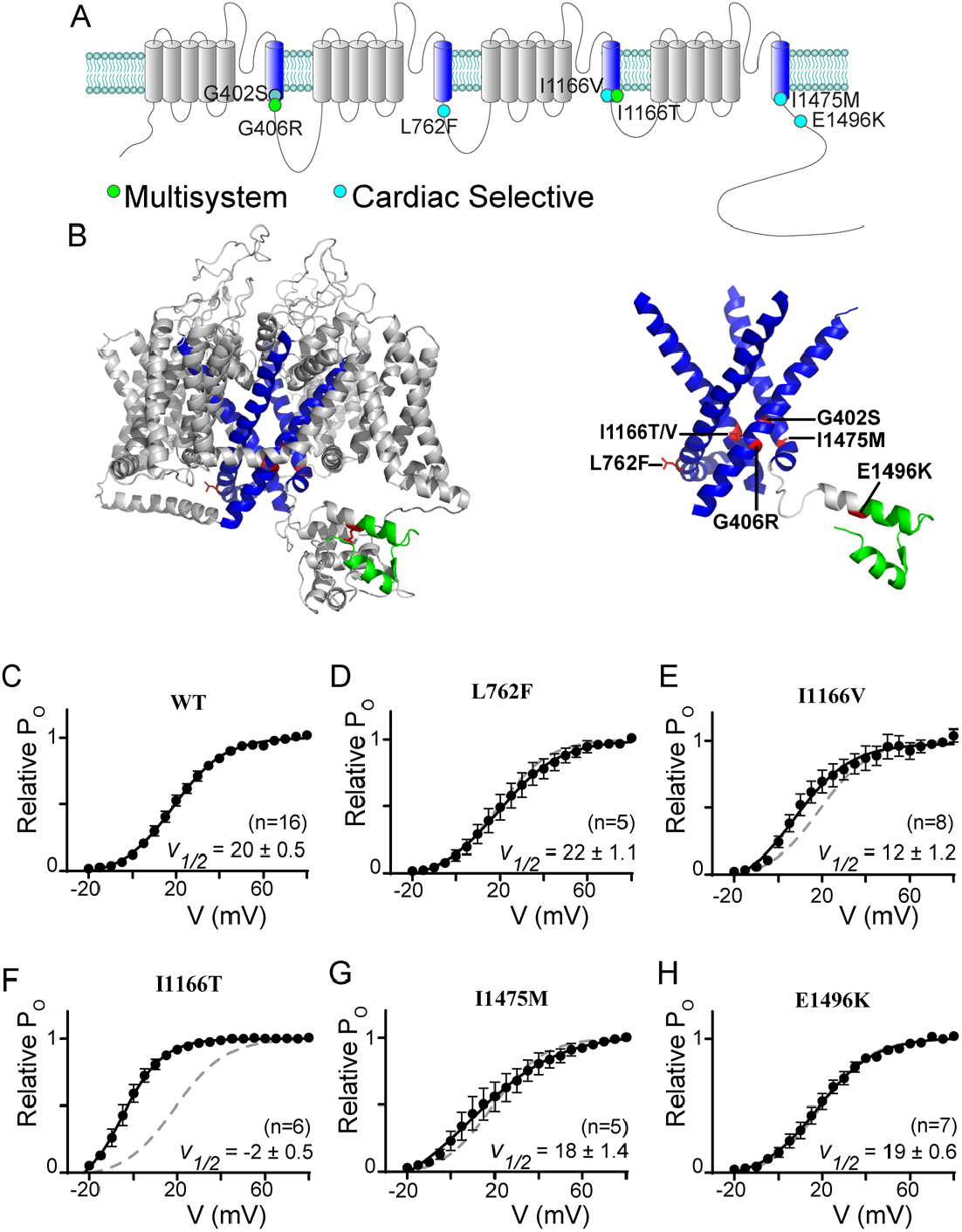
Ca_V_1.2 mutations often reside near the distal S6 region. (**A**) Cartoon depicting the location of the canonical TS mutations along with the mutations described in this study. The S6 helices are in blue; the mutations associated with a multisystem disorder including neurological symptoms are in green; and the cardiac selective mutations are in cyan. Note that the G402S mutation had been associated with a multisystem disorder, but the neurological effects have been attributed to hypoxic injury secondary to the cardiac effects (54), we therefore shade this mutation to note the distinction. (**B**) Homology model of Ca_V_1.2 demonstrating the locus of the mutations relative to the S6 helices shown in blue. Mutations are highlighted in red, EF-hand in green. *Right*: S1-5 helices are removed from the structural model to facilitate visualization of the mutations near the activation gate of the channel. (**C**) Activation curves determined by a standard tail current protocol (22). For all mutants WT data is reproduced as the dashed curve. Data is plotted ± SEM. V½ values were determined by a Boltzmann fit and are displayed ± standard error.

To fully appreciate the location of the mutations on the channel, we generated a homology model of Ca_V_1.2 based on the structure of Ca_V_1.1 (29, 30), enabling visualization of the mutation loci and surrounding residues (**Fig 1b**). The S6 regions are highlighted in blue, and visualization of the model structure with only the S6 regions displayed clearly demonstrates the location of the mutations near the helical-bundle that forms the channel activation gate, with the exception of the E1496K mutation which resides within the EF hand highlighted in green. Interestingly, while mutation L762F is predicted to lie just distal to the IIIS6 based on sequence, the homology model demonstrates that it does indeed reside on the S6 helix, which is extended in this domain.

Given the location of these mutations, we expect many may have effects on channel activation, as we previously showed for G406R and G402S (22). We therefore utilized a tail-current protocol (22) to define the relative activation curve for each mutation (**Fig. 1C-H**). Interestingly, despite its locus on the IIS6 helix, L762F did not significantly change the V_1/2_ of activation (**Fig. 1D**). However, when we explored the effect of I1166V, we found a small left shift in channel activation. Evaluation of I1166T revealed a much larger left shift, demonstrating the critical nature of this locus in channel activation. Next, we measured the effects of I1475M and E1496K and found that neither caused a change in the V_1/2_ of activation. Thus, only two of the S6 mutations investigated directly impact channel activation.

### Ca_V_1.2 mutations have selective effects on channel inactivation

Both canonical TS mutations have significant effects on channel inactivation, nearly eliminating VDI and blunting CDI (22). These effects have each been shown to have a clear impact on the cardiac action potential consistent with the LQTS phenotype of the patients (19, 22, 31). We therefore considered the impact of each of the mutations in this study on VDI and CDI. We begin by looking at VDI, which can be quantified through the use of a beta subunit permissive of VDI, such as β_1b_ (32), with Ba^2+^ as the charge carrier so as to remove the confounding effect of Ca^2+^ dependent processes such as CDI. Under these conditions, VDI can be viewed as the decay in current during a 300 ms step depolarization (**Fig. 2A**, blue shade). Quantification of VDI is given by the metric *r*_300_, which is the ratio of current remaining after 300 ms as compared to peak current such that a value of 1 would equate to no VDI, while 0 would indicate complete inactivation. Plotting the *r*_300_ for WT Ca_V_1.2 demonstrates the expected increase in VDI as a function of voltage (**Fig. 2A**, right). We next considered whether a VDI deficit may underlie the phenotype of patients harboring one of the mutations which lacked an activation shift. Indeed, evaluation of L762F demonstrated a near complete loss of VDI (**Fig. 2B**). Looking at E1496K (**Fig. 2C**), we saw a significant decrease in VDI, although to a lesser extent as compared to L762F. However, when we evaluated I1475M we found that VDI was preserved, leaving the mechanism underlying the patient phenotype for this mutation unresolved. Finally, we evaluated the mutations I1166V and I1166T, which altered channel activation; however, VDI remained unperturbed compared to WT.

**Figure 2.**
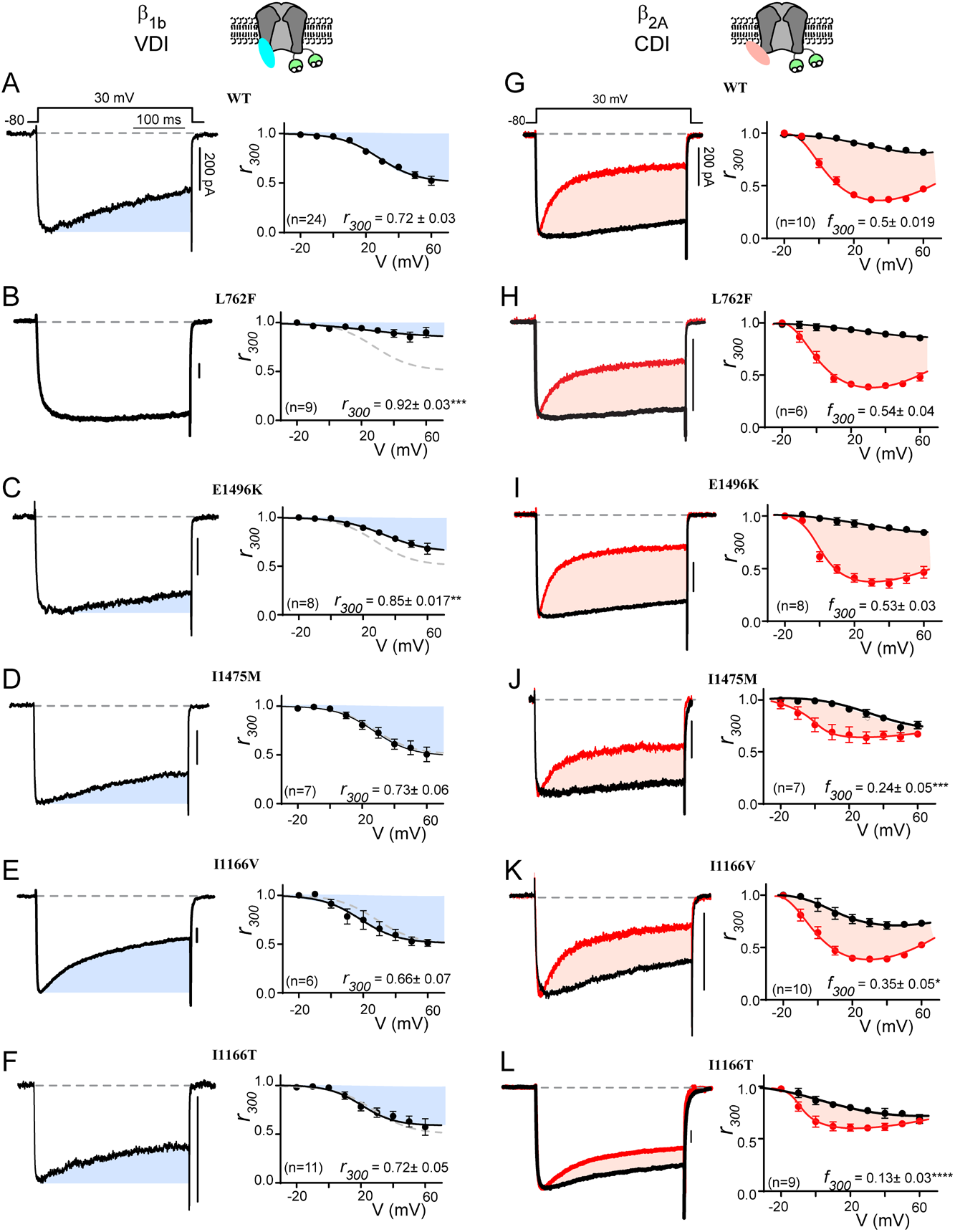
Ca_V_1.2 mutations selectively alter CDI vs. VDI. (**A**) Exemplar whole cell current traces (*left*) in Ba^2+^ demonstrate robust VDI (shaded blue) in WT Ca_V_1.2 expressed with the β_1b_ auxiliary subunit. Population data plotted for multiple voltages (*right*) demonstrates an increase in VDI with increasing voltage. Blue shading indicates the amount of VDI. *r*_300_ quantifies VDI as the ratio of current remaining after 300 ms and is displayed for the 30 mV step. Data is plotted ± SEM here and throughout. (**B, C**) VDI is significantly decreased by mutations L762F and E1496K (**D-F**) Mutations I1475M, I1166V and I1166T (respectively) do not alter VDI. (**G**) Exemplar whole cell current traces (*left*) in Ba^2+^ (black) and Ca^2+^ (red) demonstrate robust CDI (shaded peach) in WT Ca_V_1.2 expressed with the β_2A_ auxiliary subunit. Population data plotting the *r*_300_ value for multiple voltages (*right*) demonstrates minimal VDI (black) under these experimental conditions, and an expected U-shaped dependence of CDI on voltage (red). Peach shading indicates the amount of CDI, quantified by the difference between *r*_300_ values in Ca^2+^ vs. Ba^2+^ (*f*_300_). The *f*_300_ value is given for the 30 mV data and is displayed ± SEM. Error bars indicate SEM here and throughout. (**H, I**) CDI is not significantly altered by mutations L762F or E1496K. (**J-L**) Mutations I1475M, I1166V and I1166T (respectively) reduce CDI, with I1166T having the largest effect. (*p ≤ 0.05, **p ≤ 0.01, ***p ≤ 0.001, ****p ≤ 0.0001, determined via a one-way ANOVA with Dunnett’s multiple comparison test)

We next considered the effect of each mutation on CDI. To do this, we utilized the β_2A_ subunit which minimizes VDI (32), thus enhancing resolution of CDI in relative isolation. This decrease in VDI can be seen in the Ba^2+^ current trace (**Fig. 2G**, black), where inactivation is minimal. However, when Ca^2+^ is now used as the charge carrier through the channel (**Fig. 2G**, red), robust inactivation can be seen and CDI can be defined as the difference between the inactivation in Ca^2+^ as compared to Ba^2+^ (peach shaded area). Again, the metric *r*_300_ quantifies the extent of inactivation in Ca^2+^ (red) and Ba^2+^ (black), while *f*_300_ is quantified as the difference between the Ca^2+^ and Ba^2+^ *r*_300_ values and provides a metric of pure CDI. Repeating this measurement for the L762F and E1496K mutations demonstrated no effect on CDI (**Fig. 2H, I**), consistent with the lack of change in channel activation. However, evaluation of I1475M, which also displayed normal activation parameters, revealed a significant decrease in CDI across multiple voltages (**Fig. 2J**). Thus, this mutation deviates from previous L-type channel mutations which caused a change in CDI secondary to an activation change (22, 23). Finally, I1166V and I1166T (**Fig. 2K, L**) also displayed a significant decrease in CDI, with I1166T again displaying a more severe effect. Overall, evaluation of these mutations demonstrates more selective effects on channel regulation, with separable effects on VDI, CDI and activation.

### Reduced CDI at maximal Ca^2+^ concentrations revealed by Ca^2+^ uncaging

Our previous work has demonstrated that a reduction in CDI can occur through more than one mechanism (22). We therefore wanted to look more closely at the underlying deficits in the three CDI deficient channels. To do this, we considered an allosteric model for Ca_V_1.2 gating in which channels initially open in a Ca^2+^ free configuration with a relatively high open probability (P_O_), known as mode 1 (33). Upon Ca^2+^ binding to calmodulin (CaM,) channels transition into a low P_O_ state (mode Ca) (34), resulting in CDI. Within this scheme, CDI which is viewed at the whole cell level consists of two components: F_CDI_ is the fraction of channels that enter mode Ca and is dependent on the Ca^2+^ driving force; while CDI_Max_ is the maximum amount of CDI you would get if all channels were to enter mode Ca^2+^, and is dependent on the difference between the P_O_ of mode 1 vs. mode Ca (**Fig. 3A**) (23). Our prior work has demonstrated that pathogenic Ca_V_1.2 mutations can exert an effect on CDI by changing either of these two components (22). We therefore sought to measure the effect of the CDI-altering mutations on CDI_Max_. To do this, we utilized Ca^2+^ photo-uncaging paired with simultaneous patch clamp recordings and Ca^2+^ imaging. Within this setup, our goal was to drive CDI with a known concentration of Ca^2+^, enabling us to determine the extent of CDI as a function of Ca^2+^ concentration. DM-Nitrophen (DMNP) was used as the Ca^2+^ cage and added to our intracellular solution along with the Ca^2+^ indicator dye Flo4-FF. We could then release Ca^2+^ into the cytosol of the cell using a UV flash (**Fig. 3A**), while quantifying the amount of Ca^2+^ released through fluorescent imaging (22).

**Figure 3.**
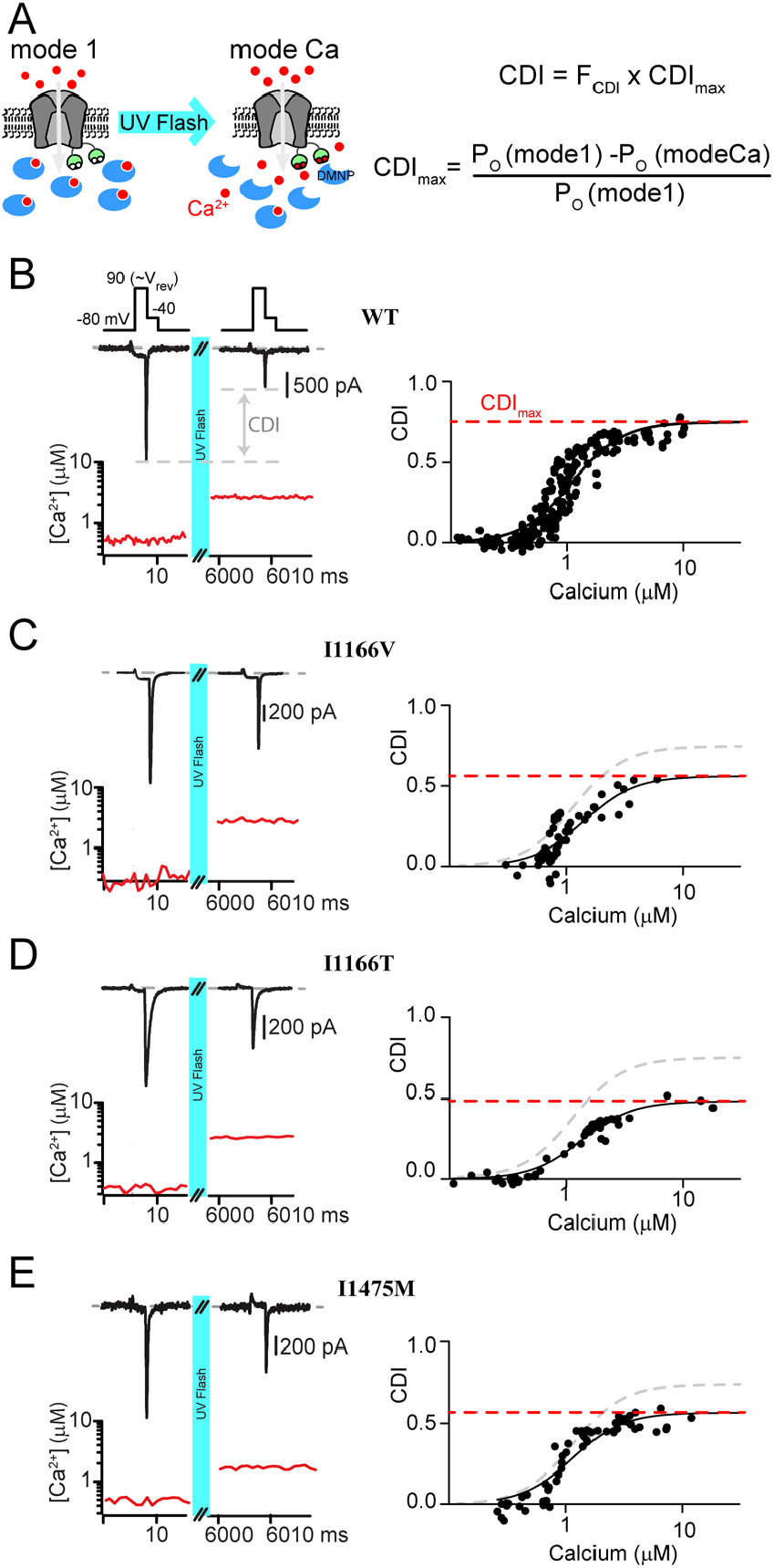
Measuring CDI as a function of Ca^2+^ concentration. (**A**) *Left:* Cartoon demonstrating the experimental setup. The Ca^2+^ cage DMNP (blue) is dialyzed into the cytosol through the patch pipette, keeping intracellular Ca^2+^ low at baseline, enabling mode 1. Application of a UV flash releases the Ca^2+^ cage, driving channels into mode Ca, which exhibits a reduced open probability. *Right:* Equations describing total CDI as a function of F_CDI_, the fraction of channels in mode C;, and CDI_Max_, the maximum amount of CDI that can be achieved if all channels were to enter mode Ca. (**B**) *Top:* Exemplar WT tail current data used to measure CDI in response to an increase in cytosolic Ca^2+^. Depolarization to the reversal potential enables channel opening without Ca^2+^ influx, followed by a step to the test potential of 40 mV, where a tail current is elicited. Tail current amplitude is significantly reduced following UV uncaging, indicative of CDI. *Bottom:* Ca^2+^ concentration is measured during the recording (red) using quantitative Ca^2+^ imaging. *Right:* Repetition of the protocol for many concentrations of Ca^2+^ resolves the CDI vs. Ca^2+^ relation. Data is fit by a Boltzmann distribution, with CDI_Max_ displayed as the dashed red line. (**C-E**) Application of the protocol to I1166V, I1166T and I1475M (respectively) demonstrates a decrease in CDI_Max_ for all mutations, with III66T having the larges effect.

Our setup is similar to our prior study in which we characterized Ca_V_1.2 CDI as a function of Ca^2+^ (22);, however, in that case we utilized a mutant Ca_V_1.2 channel to prevent intracellular Ca^2+^ release from causing pore-block(35). However, the ability to quantify Ca^2+^ dependent regulation in WT Ca^2+^ channels has since been demonstrated for Ca_V_2.1 channels (36). Given the potential for a channel mutation to impact our results, we opted to adapt this method for unaltered Ca_V_1.2 channels. In order to do this, we utilized Ca^2+^ as the charge carrier, thus bypassing the pore block which would be induced following Ca^2+^ uncaging in the presence of an alternative charge carrier (22). Next, we ensured that the cytosolic Ca^2+^ concentration would represent the Ca^2+^ concentration at the mouth of the channel by minimizing Ca^2+^ entry through the channel. Specifically, we applied a short step to the reversal potential of the channel (90 mV), which allowed channels to open without significant entry of Ca^2+^. Tail currents were then measured during a subsequent step to -40 mV; chosen so as to allow channels to fully close with kinetics enabling robust resolution of peak tail currents. A UV flash was then applied, elevating intracellular Ca^2+^, and the tail current protocol was repeated. Under these conditions, CDI was quantified as the difference in peak tail current following uncaging relative to current prior to Ca^2+^ release (**Fig. 3B**, left). Quantitative Ca^2+^ imaging then provided a readout of the cytosolic Ca^2+^ concentration for each tail current measured (**Fig. 3B**, red) and repeating the protocol for variable amounts of Ca^2+^ released enabled the generation of a relation between CDI and cytosolic Ca^2+^ (**Fig. 3B**, right). Importantly, repeated application of the tail current protocol produced stable current amplitudes and did not alter the measured cytosolic Ca^2+^ concentration. For WT channels, we obtained a CDI curve with a K_D_= 1.1 ± 0.04 µM, Hill coefficient = 2 and CDI_max_ = 0.75 ± 0.02; a result nearly identical to our prior measurements with the mutant channel (22).

We next applied our Ca^2+^ uncaging protocol to the channels harboring CDI-altering mutations. Our previous work has demonstrated that a left shift in channel activation can be causative of a decrease in CDI_max_ (22). We therefore began by considering whether the modest left shift in activation due to the I1166V mutation also caused a reduction in CDI_max_. Indeed, application of our tail current protocol to channels harboring the I1166V mutation (**Fig. 3C**) demonstrated a significant decrease in CDI_max_ (K_D_ = 1.4 ± 0.15, Hill coefficient = 2 and CDI_max_ = 0.56 ± 0.05). We next considered the effect of the I1166T mutation, which produced a large left shift in channel activation. Likewise, this mutation resulted in a robust decrease in CDI_max_ (K_D_ = 1.27 ± 0.08, Hill coefficient = 2 and CDI_max_ = 0.48 ± 0.02). Finally, we considered the effect of the I1475M mutation, which had no shift in channel activation. Despite this lack of activation change, we found that this mutation also caused a decrease in CDI_max_ (K_D_ = 1.1 ± 0.09, Hill coefficient = 2 and CDI_max_ = 0.57 ± 0.02). Given the lack of shift in the activation curve for this mutation, this result indicates that the loss of CDI proceeds from a distinct mechanism not previously described for Ca_V_1.2 mutations.

### Single channel properties reveal a mode-switch in Ca_V_1.2 mutant channels

All CDI-altering mutations in this study demonstrated a decrease in CDI_max_ (**Fig. 3**), a metric dependent on the open probability of the channel (37). F_CDI_, on the other hand, cannot be directly measured but is determined by the Ca^2+^ entering the channel during each opening. It is therefore dependent on single-channel parameters including channel conductance and open probability. Therefore, to fully understand the mechanism underlying the CDI deficit we undertook single channel recordings for each CDI-altering mutation. For these experiments, Ba^2+^ was used as the charge carrier and channels were expressed with the β_2A_ subunit in order to facilitate the evaluation of single channel properties in the absence of inactivation (32). Application of a voltage ramp across a single Ca_V_1.2 channel resulted in channel openings with a conductance of 0.018 pA mV^-1^ (**Fig. 4A**), as previously recorded for this channel under these experimental conditions (22). Averaging numerous sweeps across multiple cells enabled the resolution of a P_O_ curve (**Fig. 4B**, black) which could be well fit by a Boltzmann distribution (red). Moreover, we measured the individual open durations during the ramp and found that channels consistently exhibited brief openings, previously described as mode 1 gating (**Fig. 4C**) (33). After collecting hundreds of sweeps, we added Bay K 8644 which is known to induce mode 2 gating (**Fig. 4A**, gray box), in which channels open at more negative potentials and with significantly longer open durations. This maneuver ensures accuracy when counting the number of channels within each patch.

**Figure 4.**
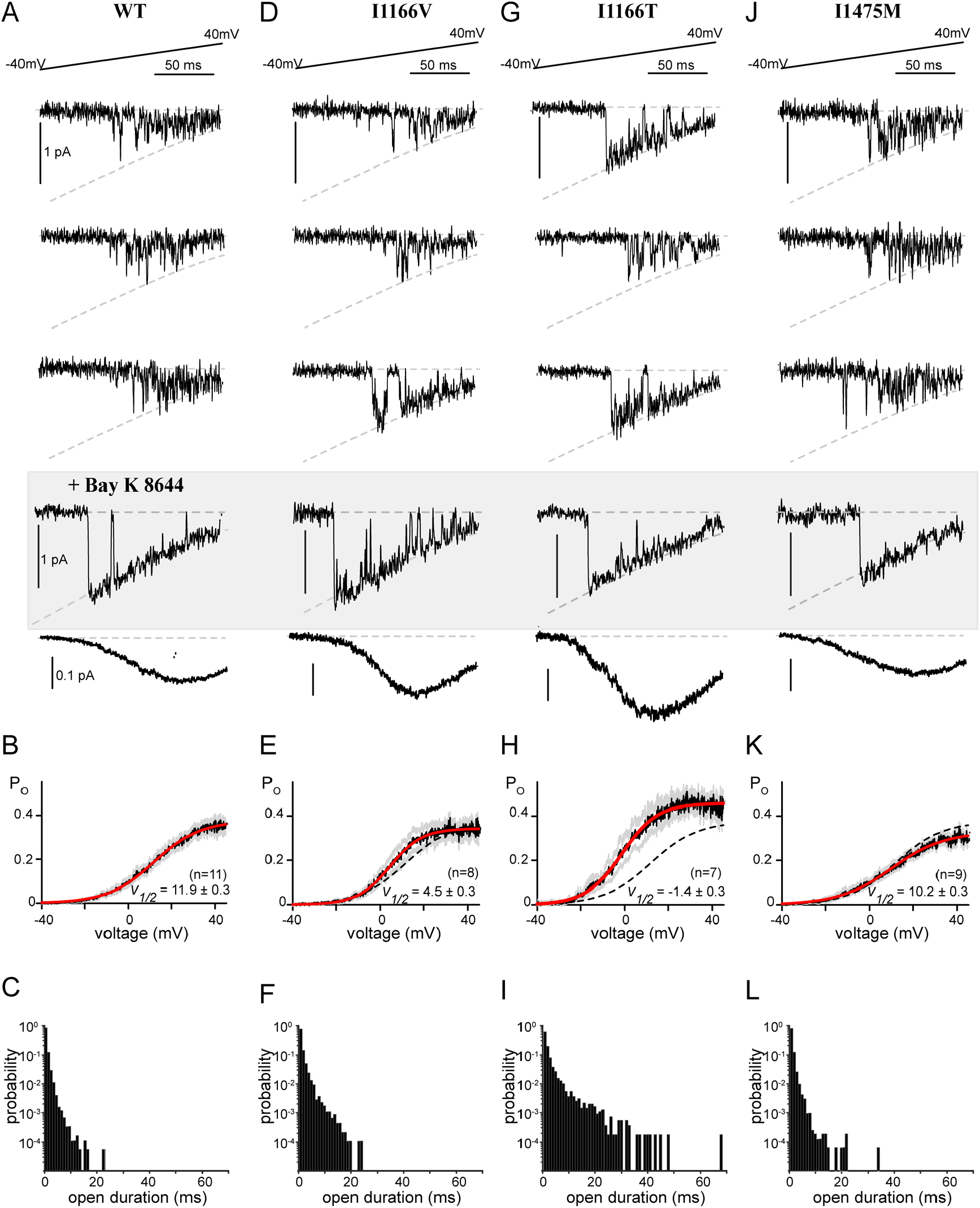
Single channel recordings reveal altered gating. (**A**) *Top:* Exemplar single channel traces in response to a voltage ramp. The ramp was applied from a voltage of -80 mV to 70 mV, however data is displayed from -40 mV to 40 mV for visualization of the single channel properties. Channel openings are seen as the downward deflections, with the dashed line indicating the single channel current as a function of voltage, as defined by the GHK equation. *Middle shaded box*: Bay K8644 was added at the conclusion of the experiment to make it easier to count the number of channels in the patch. *Bottom:* Averaging numerous sweeps generates the average current-voltage relationship for each cell. (**B**) Quantification of the open probability as a function of voltage. Black indicates the average open probability from multiple cells, gray displays the SEM and red displays the Boltzmann fit to the final data. V_1/2_ is displayed as ± the 95% confidence interval of the fit as determined by Prism (GraphPad). (**C**) Open duration histograms demonstrating the probability of opening events lasting for the indicated duration. (**D**) Exemplar single channel traces for the I1166V mutation demonstrate no change in channel conductance (dashed curve). However, while the majority of traces exhibited mode 1 gating (top 2 traces), cells would periodically display mode 2 gating (compare bottom exemplar with the Bay K 8644 exemplars). (**E**) Consistent with brief sojourns into mode 2; the average open probability as a function of voltage was somewhat left shifted as compared to WT (dashed line). Format as in **B**. (**F**) The brief periods of mode 2 gating can be seen as an increased probability of longer open durations. (**G**) Exemplar traces demonstrate significant mode 2 behavior for channels containing I1166T. (**H**) The enhanced mode 2 gating results in a large left shift in the open probability curve (red) as compared to WT (dashed line), and an overall increase in P_O_ across voltages. Format as in **B**. (**I**) Open duration histograms demonstrate a significant proportion of mode 2 openings, with durations extending beyond 40 ms. (**J-L**) No change in single channel parameters were identified in channels harboring I1475M.

We next applied the ramp protocol to channels harboring the I1166V mutation, which produced a modest left shift in channel activation in whole cell experiments (**Fig. 1**). Indeed, single channel measurements recapitulated the left shift in activation (**Fig. 4D, E**), without significant overall changes to conductance or the maximal P_O_ (P_O,max_). Interestingly, the left shift in activation was primarily due to periodic entry into mode 2. While WT Ca_V_1.2 channels will occasionally enter mode 2 without perturbation, generally application of an antagonist such as BayK 8644 (**Fig. 4**, gray box) is required to cause mode 2 gating. The mode 2 gating caused by I1166V is demonstrated in the exemplar traces displayed (**Fig. 4D**), where sweeps from the same recording switch from a mode 1 characteristic (top 2 traces) to mode 2 (bottom trace). This is further quantified by the open duration histogram (**Fig. 4F**), which displays an enhanced number of long duration openings. This mode 2 switching was even more significant when we looked at the I1166T mutation (**Fig. 4G**). There was again no change in channel conductance, however channels harboring the I1166T mutation spent a significant amount of time in mode 2, resulting in a large left shift in channel activation as well as an overall increase in P_O,max_ (**Fig. 4H**). Open duration histograms demonstrate a considerable shift to long duration openings, with durations extending beyond 40 ms (**Fig. 4I**). This remarkable behavior results in a substantial increase in channel opening across multiple voltages and correlates with the severity of the patient phenotype. Interestingly, while this increased P_O_ explains the reduction in CDI_max_ (**Fig. 3D**), it also points to an increase in F_CDI_ at lower voltages as additional Ca^2+^ would now be available to drive channels into mode Ca. Thus, the effects of each mutation on these two parameters are unlikely to be entirely independent.

Finally, we turned our attention to the I1475M mutation, which displayed a loss of CDI through a deficit in CDI_max_ despite no change in channel activation. Surprisingly, this mutation produced minimal changes to the single channel properties, with no significant change in conductance, V_1/2_ of activation or open duration (**Fig. 4 J-L)**. While we did note a slight decrease in P_O,max_ (**Fig. 4K**), this small effect cannot account for the CDI changes produced by this mutation. It therefore appears that I1475M produces a loss of CDI through a novel mechanism.

### A closer look at the neuronal effects of Ca_V_1.2 mutations

The majority of Ca_V_1.2 mutations reported in patients, and all in this study, are associated with cardiac symptoms (27, 38). However, only a select few are associated with the severe neurological phenotypes identified among TS patients (27, 38). While alterations in CDI, VDI and activation represent a recognized mechanism for inducing LQTS (19, 31), the same cannot be said for the developmental delay or ASD seen in some Ca_V_1.2 channelopathy patients. Of the mutations evaluated in this study, only I1166T has been described as causing a neurological deficit in patients. Interestingly, this mutation also results in a dramatic increase in channel activation, both through a left shift in the voltage dependance of activation, and enhanced entry into mode 2 gating (**Fig. 4H**). We therefore wanted to probe whether these biophysical properties were sufficient to alter the response of the channels to a neuronal action potential. By expressing the channels within HEK 293 cells and applying a neuronal electrical stimulus, we observe only those effects due directly to channel gating, eliminating the contribution of downstream signaling, channel trafficking or developmental changes. In order to more closely match physiological conditions, experiments were carried out in an external solution containing 1.8 mM Ca^2+^, and an internal solution containing 1 mM EGTA which permits more physiological intracellular Ca^2+^ buffering (39). In addition, we utilized the β_1B_ subunit such that both CDI and VDI are enabled (32). Ca^2+^ currents were then recorded in response to a 40 Hz train of neuronal action potentials (APs), and the amplitude of each Ca^2+^ response was measured (**Fig. 5A**). For WT channels, the peak current in response to the repetitive APs decayed slightly over the course of the stimulus (**Fig. 5B**). However, when Ca_V_1.2 channels harboring the I1166T mutation were exposed to the same stimulus, the amplitude decay (as normalized to the first peak) was significantly increased (**Fig. 5C**), demonstrating a clear difference as compared to WT channels. The same protocol applied to I1166V, I1475M, L762F or E1496K resulted in amplitudes identical to WT channels (**Fig. 5D-G**). Thus, unlike mutations which selectively alter CDI or VDI, the large changes in channel activation of I1166T are sufficient to produce a differential response to a neuronal signal.

**Figure 5.**
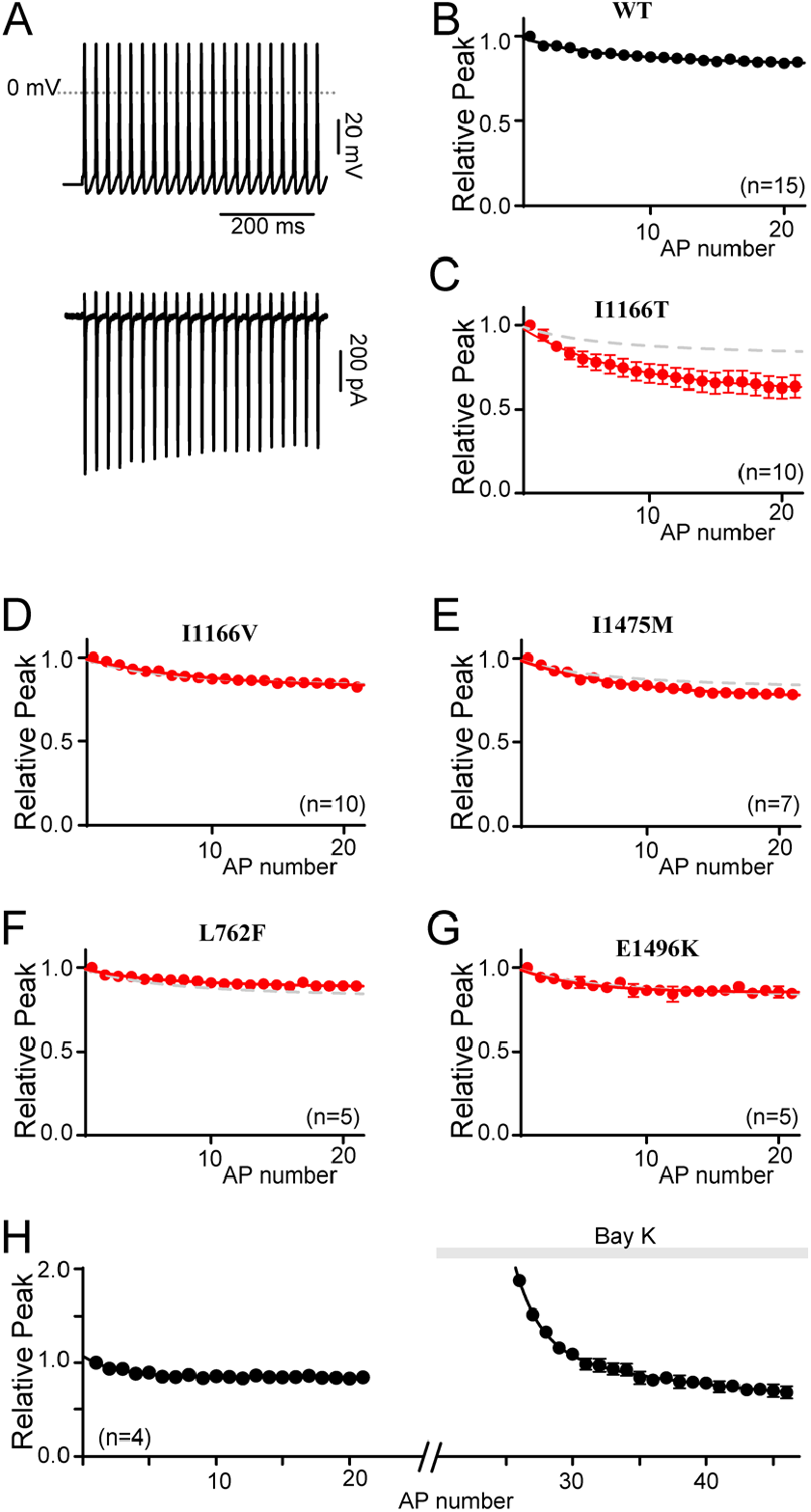
Differential responses to a neuronal stimulus. (**A**) Exemplar Ca^2+^ current response (*bottom*) to a 40 Hz train of neuronal action potentials (*top*). (**B**) Normalized peak current amplitude plotted as a function of AP number within the train reveal a modest decay in current amplitude over time. Data is displayed ± SEM here and throughout. (**C**) Normalized peak current amplitudes in channels containing I1166T (red) significantly deviate from WT (gray dashed line). (**D-G**) I1166V, I1475M, L762F and E1496K do not alter the normalized peak current amplitudes as compared to WT (gray dashed curve). (**H**) Normalized peak current amplitudes measured before and after the application of Bay K 8644 demonstrate the relative increase in current amplitude in the presence of Bay K. All currents are normalized to the first peak prior to the addition of Bay K. Data is plotted ± SEM.

While these results indicate a unique feature for the I1166T mutation, they also appear to violate the prior results which showed a loss of CDI for channels harboring the I1166T mutation. Here, I1166T appears to enhance inactivation of the current in response to a train of APs. However, under the conditions of this experiment it is unlikely that maximal CDI is achieved during the short duration of the neuronal APs, even after repetitive stimulation. Instead, we postulated that the I1166T mutation caused a large increase in the amplitude of the initial peak, however the need to normalize the data in order to compare across cells removed this component. The subsequent enhanced decay of the current through channels harboring I1166T would then result from an augmented Ca_2+_ driving force (increased F_CDI_) due to the mode 2 openings of the channel. Thus, the effects of I1166T are due to a complex interplay between a deficit in CDI_Max_ (**Fig. 3D**) and an increase in F_CDI_. To demonstrate this, we considered that the effect of the I1166T mutation is very similar to the effect of BayK 8644 when applied to WT channels. This allowed us to directly examine the impact of enhancing entry into mode 2 within the same cell. Indeed, when the AP train was applied to WT cells before and after application of Bay K 8644, a clear increase in current amplitude due to Bay K 8644 application can be seen (**Fig. 5H)**. The enhanced decay of the current then brings the amplitude back down towards the baseline. Thus, the cumulative effect of the I1166T mutation on Ca_V_1.2 current amplitude is that of a gain-of-function, as predicted by the increased activation of the mutant channel.

## Discussion

Mutations in Ca_V_1.2 have been linked to severe cardiac and neurological disorders which are often fatal in early childhood. The first two mutations described, G406R and G402S produced profound phenotypes in patients despite the fact that they occurred within a mutually exclusive exon resulting in expression in only a fraction of Ca_V_1.2 channel variants (7, 8). This severe phenotype at low expression levels may be partly due to the fact that the mutations cause significant changes to channel activation, VDI and CDI (22, 40). The mutations evaluated in the current study; however, are each expressed in a constitutive exon, resulting in much higher expression in the patient. Thus, the more selective effects on channel regulation are consistent with the patient phenotypes; severe mutations such as G406R would likely be embryonic lethal at these higher expression levels(22). Nonetheless, the gating effect of the I1166T mutation is quite severe, even in the absence of a VDI deficit. This observation fits with the relative severity of this mutation in patients, where I1166T produced the most severe phenotype of those in this study (13, 14, 16). It is also the only mutation in this study which correlates with the severe multisystem disorder described in the original TS patients harboring the G406R mutation.

Thus far, attempts to correlate biophysical changes due to Ca_V_1.2 mutations and patient phenotype have not yielded definitive results (14). This may be partly due to the comparison of experimental results across multiple conditions from different labs, or from the limited number of in-depth mechanistic studies which distinguish between different forms of channel regulation. It has also been suggested that neuronal effects of some Ca_V_1.2 mutations may be independent of Ca^2+^ permeation (41). We therefore sought to elucidate the detailed changes in channel gating due to Ca_V_1.2 mutations. We evaluated several mutations at both the whole-cell level (with conditions tuned for robust measurements of VDI vs. CDI), and further examined the mechanism underlying the CDI-altering mutations through quantitative Ca^2+^ imaging and single channel recordings. Within these parameters, we do indeed find that only the multisystem I1166T mutation produces a large increase in channel activation. Moreover, our results indicate that the gating changes of this mutation are sufficient to reproduce a change in neurological response even in the absence of downstream signaling or neuron-specific regulation (**Fig. 5**). Interestingly, both G406R and I1166T share these features of enhanced channel activation and neurological phenotypes. On the other hand, CDI and VDI deficits are present in multiple cardiac-selective mutations which did not produce an overt change in current response to a neuronal stimulus. We therefore postulate that mutations which decrease CDI or VDI, or enhance channel activation are causative of LQTS, however, only those with significant enhancement of channel activation are likely to result in ASD or neurodevelopmental disorders (**Fig. 6**). Interestingly, this result is consistent with mistuning of a voltage-dependent conformational change which has been previously identified as important for the pathogenesis of the canonical TS mutations (42); thus, in a neuron the full impact of the TS mutations may proceed from more than one pathogenic mechanism, with enhanced activation as a common feature (27).

**Figure 6.**
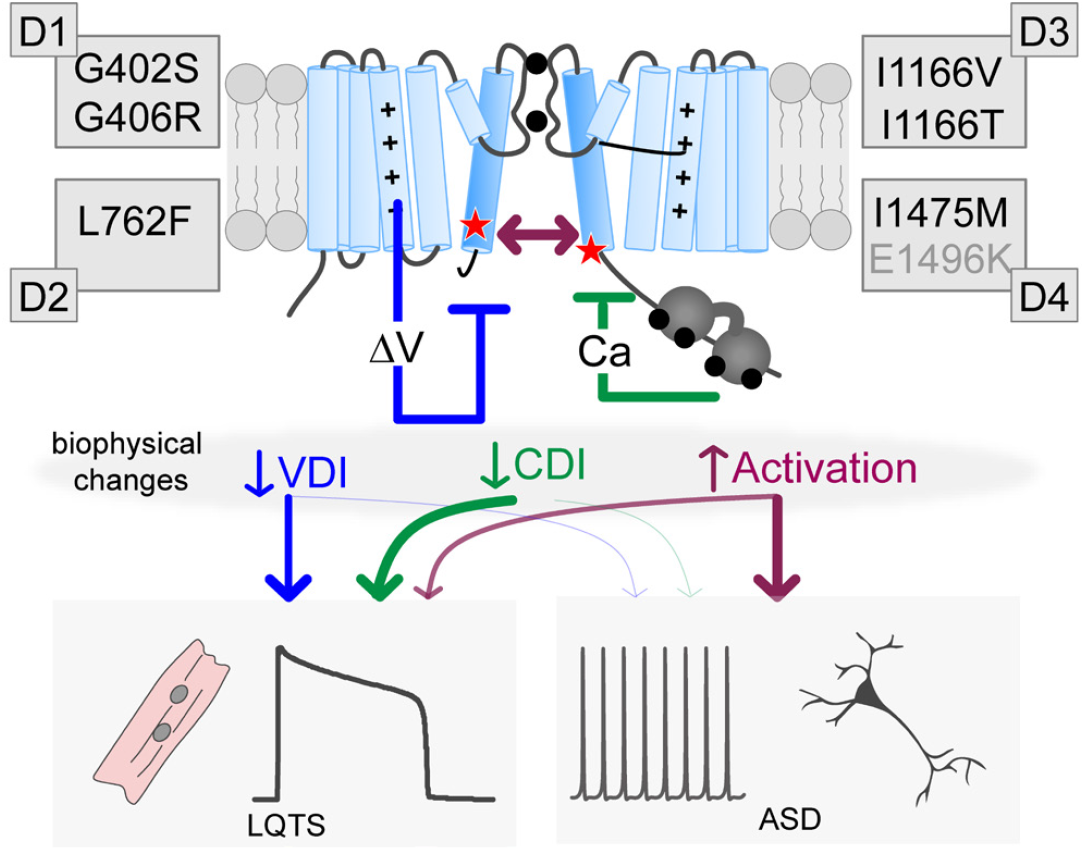
Distinct biophysical changes yield divergent pathological outcomes. Cartoon depicting the potential impact of the mutations. Each mutation can impact VDI (blue), CDI (green) and/or activation (purple), leading to LQTS. However, only large increases in channel activation appear to contribute to neurodevelopmental deficits. Note the E1496K mutation is labeled in gray as it resides outside the S6 region, nonetheless, it fits within this scheme as a VDI altering mutation.

In addition to providing insight into the mechanism underlying the pathogenesis of Ca_V_1.2 mutations, this study also expands our knowledge of the role of the S6 region in channel gating. While pathogenic mutations in Ca_V_1.2 have been identified across multiple distinct channel regions, many appear to cluster near a distal S6 and exert their impact through a change in channel gating. Interestingly, this S6 pattern is emerging as a common theme across multiple ion channels (7, 43-46), making S6-opathies an important and growing class of ion channel diseases (43). All but one of the mutations in this study occurs within a distal S6 region, and our results demonstrate that these S6 mutations can exert independent and separable effects on VDI, CDI and channel activation. The separable effects on VDI are consistent with previous models of VDI which include the distal S6 region as a locus for binding of a ‘hinged-lid’ (23). More surprising; however, is the selective effect of I1475M on CDI. Prior work has suggested that the impact of S6 mutations on CDI is often secondary to a change in the activation energy required to open the channel (23, 47). Here, we demonstrate that this is not always the case, and the S6 may participate in transducing CDI more directly. Interestingly, this effect was only seen in the distal IVS6 region, which connects the transmembrane region to the C-tail of the channel, where the Ca^2+^ sensor CaM resides on the channel. As such, it may be that the I1475M mutation disrupts the transduction of CDI following Ca^2+^ binding to CaM.

While the lack of changes in channel activation is remarkable for I1475M, the opposite is true for I1166T. For this mutation, a dramatic increase in mode 2 gating was apparent in the single channel recordings, resulting in a significant left shift in channel activation, and increased open probability across voltages (**Fig. 4G-I**). Previous studies have also suggested increased mode 2 gating in the context of a TS mutation. Erxleben et. al. showed an increase in mode 2 gating in Ca_V_1.2 channels harboring the G406R mutation; however, in this case the entry into mode 2 was thought to be controlled by an aberrant CaMKII phosphorylation site (48). However, insertion of the I1166T mutation does not form a consensus CaMKII sequence, and insertion of a V at the same site also increased the propensity for mode 2 gating, making it unlikely this mutation represents a phosphorylopathy. However, it is interesting to note that I1166 resides within a ring of hydrophobic S6 residues which are believed to stabilize the closed state of the channel (24). Mutation of the hydrophobic ‘I’ to a neutral ‘T’ residue may play a role in the enhanced opening of channels harboring the I1166T mutation. The far lesser effect of the I1166V mutation may then reflect the lack of a significant change in hydrophobicity, but nonetheless points to the importance of the I1166 residue in stabilizing the closed configuration as this mutation also produced a left shift and enhanced mode 2 gating, albeit to a far lesser extent. Likewise, the lack of an activation shift for the I1475M mutation, which resides within the IVS6 hydrophobic region, may fit with the preservation of the hydrophobicity of this residue.

It should be noted that whole cell recordings have previously been carried out on the mutations described in this study (13, 14, 16), sometimes with results which are inconsistent with the current study. In particular, E1496K and I1475M were previously reported to produce a left shift in channel activation (13). While it is not clear why these results differ, experimental conditions were quite different in that prior experiments were carried out in oocytes and activation curves were determined by a fit to the current-voltage relation. Given the important implications of the lack of activation shift for I1475M, our results from tail current protocols were confirmed by single channel recordings, demonstrating no activation shift under our experimental conditions.

Overall, this study provides an in-depth analysis of the biophysical mechanisms underlying select Ca_V_1.2 channelopathic mutations. By focusing primarily on mutations within the S6 region, this study furnishes an in depth understanding of the role of this channel segment in both activation and inactivation. Given the growing number of S6-opathy mutations being identified, such structure-function understanding may provide insight into a variety of channelopathies.

## Materials and Methods

### Molecular Biology

The α_1C_ backbone used in this study was the previously described human cardiac Ca_V_1.2 clone (22) within the pcDNA3 plasmid and corresponds to Acc# Z34810. The original plasmid used to generate this clone was a kind gift from Tuck Wah Soong (49). Channel mutations were introduced into this plasmid using the QuickChange Lightning kit from Agilent. All portions of the resulting construct that were subject to PCR were confirmed by DNA sequencing.

### Homology model

A homology model of Ca_V_1.2 was generated based on the 5GJW PDB structure of the Ca_V_1.1 channel (29). Sequences for the human Ca_V_1.2 channel used in this study and the Ca_V_1.1 channel corresponding to the PDB structure were aligned and trimmed to match and the software Modeller was used to generate the model (50). The subsequent structural model was visualized using PyMol and figures were generated from this software.

### Transfection

All patch clamp experiments were performed using HEK 293 cells transfected via the calcium phosphate method (51). Cells were transfected with the human α_1C_ subunit along with the rat α_2δ_ subunit (NM012919.2) and either rat brain β_1B_ or β_2a_ (52) as indicated. The SV40 T antigen was also co-transfected in order to enhance expression levels. To facilitate identification of transfected cells, all transfections included GFP either as a separate plasmid (for β_1B_) or as part of a GFPIR vector containing the beta subunit (β_2a_). Cells were transfected and used for patch clamp recordings within 1-3 days.

### Whole cell patch clamp

Whole cell patch clamp recordings were performed at room temperature using an Axopatch 200B amplifier (Axon Instruments). Borosilicate glass electrodes were generated with resistances between 1-3 MΩ, with series resistance compensated to >70% during the recordings. Currents were low-pass filtered at 2 kHz (4-pole Bessel filter) and sampled at 10 kHz during VDI and CDI measurements. Tail current protocols and neuronal AP recordings were recorded with 5 kHz low-pass filtering and sampled at 50 kHz so as to enable full resolution of the peak. For activation curve, CDI and VDI measurements, the internal solution contained (in mM): CsMeSO3, 114; CsCl, 5; MgATP, 4; HEPES (pH 7.4), 10; and BAPTA (1,2-bis(*o*-aminophenoxy)ethane-*N,N,N’,N’*-tetraacetic acid), 10; at 295 mOsm adjusted with CsMeSO3 and the bath solution contained (in mM): TEA-MeSO3, 102; HEPES (pH 7.4), 10; CaCl2 or BaCl2, 40; at 305 mOsm adjusted with TEA-MeSO3. Solutions for the neuronal stimulus were adjusted to more closely approximate physiological Ca^2+^ such that the internal solution contained (in mM): CsMeSO3, 114; CsCl, 5; MgATP, 4; HEPES (pH 7.4), 10; and EGTA (ethylene glycol-bis(β-aminoethyl ether)-N,N,N′,N′-tetraacetic acid), 1; at 295 mOsm adjusted with CsMeSO3 and the bath solution contained (in mM): TEA-MeSO3, 102; HEPES (pH 7.4), 40; CaCl2, 1.8; at 305 mOsm adjusted with TEA-MeSO3. Data was analyzed using custom Matlab scripts with all fits done in Prism (GraphPad). Inactivation was quantified as the ratio of current remaining after 300 ms in either Ca^2+^ or Ba^2+^ (*r*_300_), such that the difference between the *r*_300_ values recorded in Ba^2+^ versus Ca^2+^ quantified pure CDI.

### Single channel recordings

Single channel patch clamp recordings were performed at room temperature using an Axopatch 200B amplifier (Axon Instruments). Thick walled borosilicate glass electrodes were generated with resistances between 3-5 MΩ and coated with Sylgard. Currents were low-pass filtered at 2 kHz (4-pole Bessel filter) and sampled at 10 kHz. The pipette solution matched the whole cell extracellular solution and contained (in mM): TEA-MeSO3, 102; HEPES (pH 7.4), 10; BaCl2, 40; at 305 mOsm adjusted with TEA-MeSO3. The bath solution was designed to zero the membrane potential and contained (in mM): K glutamate, 132; KCl, 5; NaCl, 5; MgCl, 3; EGTA, 2, glucose, 10; and HEPES (pH 7.4), 20 at 300 mOsm adjusted with glucose.

The ramp protocol was applied from -80 mV to 70 mV over a duration of 200 ms. The leak for each sweep was manually fit using a combination of a linear and exponential functions. The unitary current amplitude was fit with the GHK equation (53), providing a readout for conductance. The voltage parameter V_s_ was allowed to vary by ± 3 mV between patches to allow for recording variability. For each cell, traces were averaged, excluding blank traces, to produce a current-voltage relation. These curves were then averaged together for different patches and the P_O_ was determined using the GHK relation. Prior to the conclusion of the experiment Bay K 8644 was added to a final concentration of ∼5 µM to allow for easy counting of the number of channels in the patch. We found that our estimate of the number of channels was consistent with the count in Bay K, therefore recordings in which the cell died prior to Bay K addition (but with >50 sweeps recorded) did not need to be excluded from the average. Patches with 1-3 channels were included in the average, and blanks were excluded such that the data corresponds to active sweeps only, as previously described for Ca_V_1.2 channels harboring TS mutations (22).

### Ca^2+^ uncaging

For simultaneous Ca^2+^ uncaging and patch clamp recordings, we used a calibrated mix of 5 μM Fluo-2 (TefLabs, Austin TX); 5 μM Fluo-2 Low Affinity (TefLabs) and 2.5 μM Alexa568 (Invitrogen, Waltham MA) as previously described (36). This combination enabled ratiometric recording across a range of Ca^2+^ concentrations. Stocks of this solution were premixed and calibrated prior to the experiments so that there was no variability in the ratios across experiments. In addition to these dyes, the internal solution contained, (in mM): CsCl, 135; HEPES (pH 7.4), 40; DMNP-EDTA, 1-4; Citrate; 1-20 mM; CaCl, 0.75-3.5. Citrate, DMNP and CaCl were adjusted to provide varying levels of Ca^2+^ steps, such that baseline calcium levels were below 100 nM as measured by the Ca^2+^ dyes. External solutions contained (in mM): TEA-MeSO3, 70; HEPES (pH 7.4), 10; and CaCl, 40. Data were analyzed by custom MATLAB software (Mathworks, MA) and final fits to the data were done in Prism (GraphPad).

## Acknowledgements

We thank Debora DiSilvestre and Josiah Owoyemi for dedicated technical support. We also thank John Hussey and Dr. Sara Codding for insightful comments and discussion. This project was supported by an NIH/NHLBI grant (1R01HL149926) and by an AHA postdoctoral fellowship (20POST35211127).

## References

1. P. J. Adams, T. P. Snutch, Calcium channelopathies: voltage-gated calcium channels. Subcell Biochem 45, 215–251 (2007).

2. K. S. Lee, E. Marban, R. W. Tsien, Inactivation of calcium channels in mammalian heart cells: joint dependence on membrane potential and intracellular calcium. J Physiol 364, 395–411 (1985).

3. R. D. Zuhlke, G. S. Pitt, K. Deisseroth, R. W. Tsien, H. Reuter, Calmodulin supports both inactivation and facilitation of L-type calcium channels. Nature 399, 159–162 (1999).

4. B. Z. Peterson, C. D. DeMaria, J. P. Adelman, D. T. Yue, Calmodulin is the Ca2+ sensor for Ca2+ - dependent inactivation of L-type calcium channels. Neuron 22, 549–558 (1999).

5. S. Beyl et al., Different pathways for activation and deactivation in CaV1.2: a minimal gating model. J Gen Physiol 134, 231-241; S231-232 (2009).

6. H. Liang et al., Unified mechanisms of Ca2+ regulation across the Ca2+ channel family. Neuron 39, 951–960 (2003).

7. I. Splawski et al., Severe arrhythmia disorder caused by cardiac L-type calcium channel mutations. Proc Natl Acad Sci U S A 102, 8089–8096; discussion 8086-8088 (2005).

8. I. Splawski et al., Ca(V)1.2 calcium channel dysfunction causes a multisystem disorder including arrhythmia and autism. Cell 119, 19–31 (2004).

9. C. Napolitano, I. Splawski, K. W. Timothy, R. Bloise, S. G. Priori, “Timothy Syndrome” in GeneReviews(R), R.A. Pagon et al., Eds. (Seattle (WA), 1993).

10. M. L. Marks, D. L. Trippel, M. T. Keating, Long QT syndrome associated with syndactyly identified in females. The American journal of cardiology 76, 744–745 (1995).

11. M. L. Marks, S. L. Whisler, C. Clericuzio, M. Keating, A new form of long QT syndrome associated with syndactyly. Journal of the American College of Cardiology 25, 59–64 (1995).

12. A. T. Lu, X. Dai, J. A. Martinez-Agosto, R. M. Cantor, Support for calcium channel gene defects in autism spectrum disorders. Mol Autism 3, 18 (2012).

13. K. Wemhoner et al., Gain-of-function mutations in the calcium channel CACNA1C (Cav1.2) cause non-syndromic long-QT but not Timothy syndrome. J Mol Cell Cardiol 80, 186–195 (2015).

14. A. P. Landstrom et al., Novel long QT syndrome-associated missense mutation, L762F, in CACNA1C-encoded L-type calcium channel imparts a slower inactivation tau and increased sustained and window current. International journal of cardiology 220, 290–298 (2016).

15. J. Gillis et al., Long QT, syndactyly, joint contractures, stroke and novel CACNA1C mutation: expanding the spectrum of Timothy syndrome. American journal of medical genetics. Part A 158A, 182–187 (2012).

16. N. J. Boczek et al., Novel Timothy syndrome mutation leading to increase in CACNA1C window current. Heart Rhythm 12, 211–219 (2015).

17. B. A. Alseikhan, C. D. DeMaria, H. M. Colecraft, D. T. Yue, Engineered calmodulins reveal the unexpected eminence of Ca2+ channel inactivation in controlling heart excitation. Proc Natl Acad Sci U S A 99, 17185–17190 (2002).

18. A. L. George, Jr., Calmodulinopathy: a genetic trilogy. Heart Rhythm 12, 423–424 (2015).

19. D. Morales, T. Hermosilla, D. Varela, Calcium-dependent inactivation controls cardiac L-type Ca(2+) currents under beta-adrenergic stimulation. J Gen Physiol 151, 786–797 (2019).

20. J. Ozawa et al., A novel CACNA1C mutation identified in a patient with Timothy syndrome without syndactyly exerts both marked loss- and gain-of-function effects. HeartRhythm Case Rep 4, 273–277 (2018).

21. P. Liao, T. W. Soong, CaV1.2 channelopathies: from arrhythmias to autism, bipolar disorder, and immunodeficiency. Pflugers Arch 460, 353–359 (2010).

22. I. E. Dick, R. Joshi-Mukherjee, W. Yang, D. T. Yue, Arrhythmogenesis in Timothy Syndrome is associated with defects in Ca(2+)-dependent inactivation. Nat Commun 7, 10370 (2016).

23. M. R. Tadross, M. Ben Johny, D. T. Yue,Molecular endpoints of Ca2+/calmodulin-and voltage-dependent inactivation of Ca(v)1.3 channels. J Gen Physiol 135, 197–215 (2010).

24. S. Hering et al., Calcium channel gating. Pflugers Arch 470, 1291–1309 (2018).

25. M. Kudrnac et al., Coupled and independent contributions of residues in IS6 and IIS6 to activation gating of CaV1.2. J Biol Chem 284, 12276–12284 (2009).

26. A. Raybaud et al., The role of the GX9GX3G motif in the gating of high voltage-activated Ca2+ channels. J Biol Chem 281, 39424–39436 (2006).

27. A. Marcantoni, C. Calorio, E. Hidisoglu, G. Chiantia, E. Carbone, Cav1.2 channelopathies causing autism: new hallmarks on Timothy syndrome. Pflugers Arch 472, 775–789 (2020).

28. J. Kim, S. Ghosh, D. A. Nunziato, G. S. Pitt, Identification of the components controlling inactivation of voltage-gated Ca2+ channels. Neuron 41, 745–754 (2004).

29. J. Wu et al., Structure of the voltage-gated calcium channel Ca(v)1.1 at 3.6 A resolution. Nature 537, 191–196 (2016).

30. Y. Zhao et al., Molecular Basis for Ligand Modulation of a Mammalian Voltage-Gated Ca(2+) Channel. Cell 177, 1495–1506 e1412 (2019).

31. S. Morotti, E. Grandi, A. Summa, K. S. Ginsburg, D. M. Bers, Theoretical study of L-type Ca(2+) current inactivation kinetics during action potential repolarization and early afterdepolarizations. J Physiol 590, 4465–4481 (2012).

32. S. K. Wei et al., Ca(2+) channel modulation by recombinant auxiliary beta subunits expressed in young adult heart cells. Circ Res 86, 175–184 (2000).

33. P. Hess, J. B. Lansman, R. W. Tsien, Different modes of Ca channel gating behaviour favoured by dihydropyridine Ca agonists and antagonists. Nature 311, 538–544 (1984).

34. J. P. Imredy, D. T. Yue, Mechanism of Ca(2+)-sensitive inactivation of L-type Ca2+ channels. Neuron 12, 1301–1318 (1994).

35. R. K. Cloues, S. M. Cibulsky, W. A. Sather, Ion interactions in the high-affinity binding locus of a voltage-gated Ca(2+) channel. J Gen Physiol 116, 569–586 (2000).

36. S. R. Lee, P. J. Adams, D. T. Yue, Large Ca(2)(+)-dependent facilitation of Ca(V)2.1 channels revealed by Ca(2)(+) photo-uncaging. J Physiol 593, 2753–2778 (2015).

37. M. R. Tadross, D. T. Yue, Systematic mapping of the state dependence of voltage-and Ca2+-dependent inactivation using simple open-channel measurements. J Gen Physiol (2010).

38. N. J. Boczek et al., Identification and Functional Characterization of a Novel CACNA1C-Mediated Cardiac Disorder Characterized by Prolonged QT Intervals With Hypertrophic Cardiomyopathy, Congenital Heart Defects, and Sudden Cardiac Death. Circulation. Arrhythmia and electrophysiology 8, 1122–1132 (2015).

39. M. D. Stern, Buffering of calcium in the vicinity of a channel pore. Cell calcium 13, 183–192 (1992).

40. C. F. Barrett, R. W. Tsien, The Timothy syndrome mutation differentially affects voltage- and calcium-dependent inactivation of CaV1.2 L-type calcium channels. Proc Natl Acad Sci U S A 105, 2157–2162 (2008).

41. J. F. Krey et al., Timothy syndrome is associated with activity-dependent dendritic retraction in rodent and human neurons. Nature neuroscience 16, 201–209 (2013).

42. B. Li, M. R. Tadross, R. W. Tsien, Sequential ionic and conformational signaling by calcium channels drives neuronal gene expression. Science 351, 863–867 (2016).

43. P. Lory, S. Nicole, A. Monteil, Neuronal Cav3 channelopathies: recent progress and perspectives. Pflugers Arch 472, 831–844 (2020).

44. M. Kschonsak et al., Structure of the human sodium leak channel NALCN. Nature 587, 313–318 (2020).

45. S. Biel et al., Mutation in S6 domain of HCN4 channel in patient with suspected Brugada syndrome modifies channel function. Pflugers Arch 468, 1663–1671 (2016).

46. C. Shelley, J. P. Whitt, J. R. Montgomery, A. L. Meredith, Phosphorylation of a constitutive serine inhibits BK channel variants containing the alternate exon “SRKR”. J Gen Physiol 142, 585–598 (2013).

47. I. E. Dick, M. R. Tadross, D. T. Yue (2010) Diverging mechanisms underlying calcium inactivation defects leading to Timothy Syndrome.in Society for Neuroscience (San Diego), p 848.811/F822.

48. C. Erxleben et al., Cyclosporin and Timothy syndrome increase mode 2 gating of CaV1.2 calcium channels through aberrant phosphorylation of S6 helices. Proc Natl Acad Sci U S A 103, 3932–3937 (2006).

49. Z. Z. Tang et al., Transcript scanning reveals novel and extensive splice variations in human l-type voltage-gated calcium channel, Cav1.2 alpha1 subunit. J Biol Chem 279, 44335–44343 (2004).

50. B. Webb, A. Sali, Comparative Protein Structure Modeling Using MODELLER. Curr Protoc Protein Sci 86, 2 9 1–2 9 37 (2016).

51. D. L. Brody, P. G. Patil, J. G. Mulle, T. P. Snutch, D. T. Yue, Bursts of action potential waveforms relieve G-protein inhibition of recombinant P/Q-type Ca2+ channels in HEK 293 cells. J Physiol 499 (Pt 3), 637–644 (1997).

52. E. Perez-Reyes et al., Cloning and expression of a cardiac/brain beta subunit of the L-type calcium channel. J.Biol.Chem. 267, 1792–1797 (1992).

53. B. Hille, Ion channels of excitable membranes (Sinauer, Sunderland, Mass., ed. 3rd, 2001), pp. xviii, 814 p.

54. S. Frohler et al., Exome sequencing helped the fine diagnosis of two siblings afflicted with atypical Timothy syndrome (TS2). BMC Med Genet 15, 48 (2014).

